# Home, head direction stability and grid cell distortion

**DOI:** 10.1101/602771

**Authors:** Juan Ignacio Sanguinetti-Scheck, Michael Brecht

**Author notes:** **Corresponding author:** Michael Brecht; Humboldt-Universität zu Berlin, Philippstr. 13, Haus 6, 10115 Berlin, Germany.

## Abstract

The home is a unique location in the life of humans and animals. Numerous behavioral studies investigating homing indicate that many animals maintain an online representation of the direction of the home, a home vector. Here we placed the rat’s home cage in the arena, while recording neurons in the animal’s parasubiculum and medial entorhinal cortex. From a pellet hoarding paradigm it became evident that the home cage induced locomotion patterns characteristic of homing behaviors. We did not observe home-vector cells. We found that head-direction signals were unaffected by home location. However, grid cells were distorted in the presence of the home cage. While they did not globally remap, single firing fields were translocated towards the home. These effects appeared to be geometrical in nature rather than a home-specific distortion. Our work suggests that medial entorhinal cortex and parasubiculum do not contain an explicit neural representation of the home direction.

## Introduction

Animals maintain and update a representation of their location in space. Such mapping abilities allow them to navigate their surroundings in search for food, safety, mates or their kin. In the case of the migratory Bar-tailed Godwit such navigation can involve a non-stop 11000 km flight from New Zealand wintering grounds to artic Siberian breeding grounds (Gill et al., 2005) a truly incredible navigational feat. The home is a unique location in the life of humans and animals. Numerous behavioral studies have researched homing; in pigeons (Alleva et al., 1975), in bees is returning to the hive (Menzel et al., 2005), in salmon returning to the stream where they were born in (Neave, 1964), in bats returning to their cave from foraging their favorite tree (Tsoar et al., 2011) and many other species. Safety considerations shape rat exploratory behaviors and lab rats naturally organize their behavior around their home-cage (Whishaw et al., 2006).

A consensus emerged that patterns of navigation show remarkable and systematic variations relative to the home location. In particular, animals often show variable and complicated trajectories outgoing from the home only to return on relatively direct, straight incoming trajectories back to the home location. Some of the best evidence for path integration in the animal kingdom comes pup retrieval experiments of the Mittelstaedts (Mittelstaedt and Mittelstaedt, 1980), where the authors demonstrated path integration by the mother’s tendency to return pups on a straight trajectory back to the home. In the proximity of the home startling stimuli will very often induce a short-latency escape maneuver straight back home. Thus, many animals maintain an immediately accessible online representation of the direction of the home, a representation we refer to as home vector. In our paper we ask how this home vector might be represented in the well-known neural substrates of navigation. Head Direction cells, Goal Direction cells and Grid cells have been hypothesized to sustain vectorial navigation (Kubie and Fenton, 2009; Valerio and Taube, 2012; Sarel et al., 2017; Banino et al., 2018).

Head Direction cells, present in several brain structures including the anterior dorsal nucleus of the thalamus, the presubiculum, the medial enthorinal cortex (MEC) and the Parasubiculum (PaS), have sharp tuning curves in relation the animals orientation in space (Taube et al., 1990; Taube, 1995; Tang et al., 2016). In lab navigation tasks the accuracy of Head Direction cells also predicts successful navigation (Valerio and Taube, 2012).

Grid cells of the MEC and parasubiculum are known to be active in multiple spatial firing fields which tile the whole environment forming a periodic hexagonal lattice (Fyhn, 2004; Hafting et al., 2005; Boccara et al., 2010; Tang et al., 2016). Even though recent work points towards differences in field firing rates as a way for a single cell to encode different environments and local positional information (Diehl et al., 2017; Ismakov et al., 2017; Stensola et al., 2015).

We were interested in the role of the parasubiculum in homing. The parasubiculum is a long (3mm) and thin (300 microns) parahippocampal cortex wrapped around the medial and dorsal boundaries of the MEC. It contains both a high proportion of head directional and spatially selective cells, including grid cells and border cells (Boccara et al., 2010), and connects selectively to pyramidal patches in layer 2 of the MEC (Tang et al., 2016). Specifically, we ask the following questions:

(1) Do rats care about their home cage?
(2) Are there home-direction cells, whose discharge is tuned to the home location?
(3) Are head-direction signals altered or distorted by the home location?
(4) Are grid cell signals altered or distorted by the home location?

Our data indicate that the home-cage induce homing-like behaviors. We did not observe home-direction cells and found that head-direction signals are not affected by the home location. Grid cell signals were locally altered by the home cage location, but the effects appeared to be more geometrical in nature rather than home-specific.

## Results

In our study we addressed the question how neurons in the rat medial entorhinal cortex and the rat parasubiculum represent the home location. To this end we recorded from freely moving male Long Evans rats (n=6) using tetrodes. Rats were familiarized for two weeks with a 1m × 1m squared environment with round edges containing a principal cue card multiple irregularly shaped sub-cues, both on the walls or the floor of the arena (Figure1A). During the same period, we housed rats in custom-modified home-cage modified with 2 side doors (Figure1A). To assess if the home cage affected the rat’s spatial behavior we studied rats performing pellet hoarding. During pellet hoarding (Figure 1; Wolfe, 1939) rats forage for food pellets and perform high-speed return vectors towards their safe location. In our setting (Figure 1A) rats foraged large food pellets in a 1 meter arena in the presence of their home cage. Without specific prior training rats foraged these pellets and cached them in their home cage. The behavior was stereotypical, consisting of high speed return trips as previously described in the literature (Maaswinkel and Whishaw, 1999; Winter et al., 2018)(Figure 1 B). Running speed was visibly lower during exploratory trips away from their home (Figure 1 C). As has been described in the literature (Tchernichovski et al., 1998; Wallace et al., 2008; Winter et al., 2018) incoming trips (Figure 1D) consisted of higher speeds (KS normality, p=0.747, ttest, p<0.001) than outgoing (Figure 1E). Thus, in the presence of large food pellets the home cage in the arena setting greatly altered the rat’s locomotion patterns and divided them into irregular, slower exploratory outgoing trajectories and relatively straight, faster return trips to the home cage. These observations suggest that the home cage can induce homing behaviors, even in a scenario where the rat is well adapted to the global environment.

**Figure 1.**
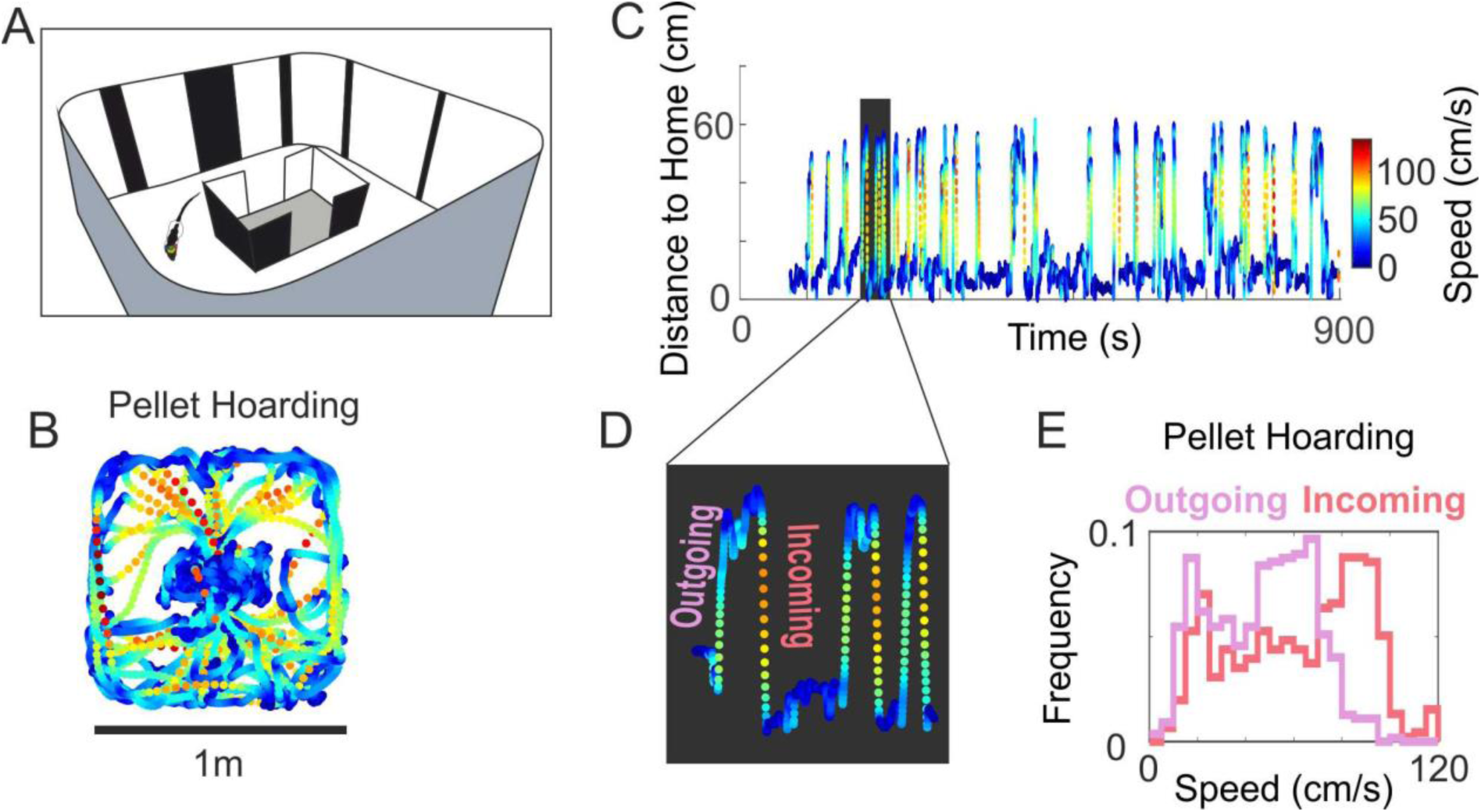
In a pellet hoarding paradigm the home cage induces differential outgoing and incoming locomotion reflecting homing behavior. A) Top: Schematics of the environment with the home cage in the center; the animal’s home cage was modified such that not only the lid could be removed, but that also two gaps in the sides of the cage could be opened. Walls are covered in complex cues. Bottom: We recorded 12-20 minute sessions without removing the rat form the arena. B) Speed profile. Trajectory of the animal in the environment are color-coded according to running speed while the rat is hoarding pellets in the home in the center of the arena. C) Rat performed exploratory trips from the home. Speed is color coded. Distance to the home plot against time for a rat performing pellet hoarding behavior. D) Magnified view of three single foraging trips in C. E) Histograms of proportions of speeds for outgoing (pink) and incoming (red) exploratory trips for example in C. (p<0.001, Krustal-Wallis). Overall median speeds of incoming trips (n= 13 sessions) were significantly higher than outgoing trips (KS normality, p=0.747, ttest, p<0.001).

We wondered how the presence of the home cage would affect head direction cells and grid cells, and whether we could find any home vector representation constantly representing the egocentric direction of the home, a home vector. To concentrate on the effect of the presence of the home, instead of differences introduced by the hoarding behavior, we studied neural activity while rats performed the same small treat random foraging behavior in the absence and presence of the home. The introduction of the home in the absence of large pellets, resulted in comparable behavior to the open field in terms of speed (not shown). Median incoming and outgoing trips from the home in the absence of pellets to be hoarded were not different (KS normality, p=0.747, ttest, p=0.9).

We recorded sessions of 12-25 minutes starting with an open field recording and following with sessions where we placed the rat’s own home-cage in different places in the arena. In order not to disturb the familiarity of the rat with the arena, we did not remove or disorient the rat between sessions. This was facilitated by the use of a wireless logger system for recording which allowed us to test exclusively for the effects of locally altering the internal geometry of the environment. Using tetrodes we recorded (n=500) cells in the parasubiculum, MEC and medial MEC (Ray et al., 2017), which we classified into Pure head direction Cells (n=90), Pure grid cells (n=35), conjunctive grid cells (n=50) and Rest (n=325). We analyzed whether the presence of the home resulted in pure or conjunctive representations of egocentric home direction and whether the home presence affects the encoding of the environment by head direction cells and grid cells.

### Head direction activity is not affected by home cage presence

As the first step of our analysis of neural responses we assessed how the home cage affected head direction signals (Figure 2). We recorded (n=68) head direction cells in the PaS/MEC with the presence of the home in the center of the environment or the home rotated and translated to the edge of the environment (n=20; Figure 2A) and found that pure head direction cells remained stable during both the introduction and the translation of the animals home (Figure 2B/C). Differences in Rayleigh vector length of head direction in comparison to the open field condition were centered on zero showing no bias towards an increase or decrease of head directional coding (Figure 2D). Head direction cells also maintained their angular preference. This is obvious from the cumulative distribution of angle differences between home cage and open field condition, which is narrowly centered at 0 degrees (Figure 2E). Overall, there was a strong correlation between the angular rate distributions with and without the home (Figure2F). The head directionality of conjunctive grid cells was also unaffected by the presence of the home (Figure 2G,H), both in angle preference (Figure 2H) and vector length (not shown, KS Normality Test, p=0.036, Signed-Rank Test, p=0.856, N=39). In a subset head directional cells (n=20, Head Direction and Conjuntive Grids cells), we studied head directional encoding while the rat performed hoarding behavior in the presence of the home, we found head directionality to remain stable with the performance of the task (not shown). All in all we conclude that the presence of the home does not affect head direction tuning properties.

**Figure 2.**
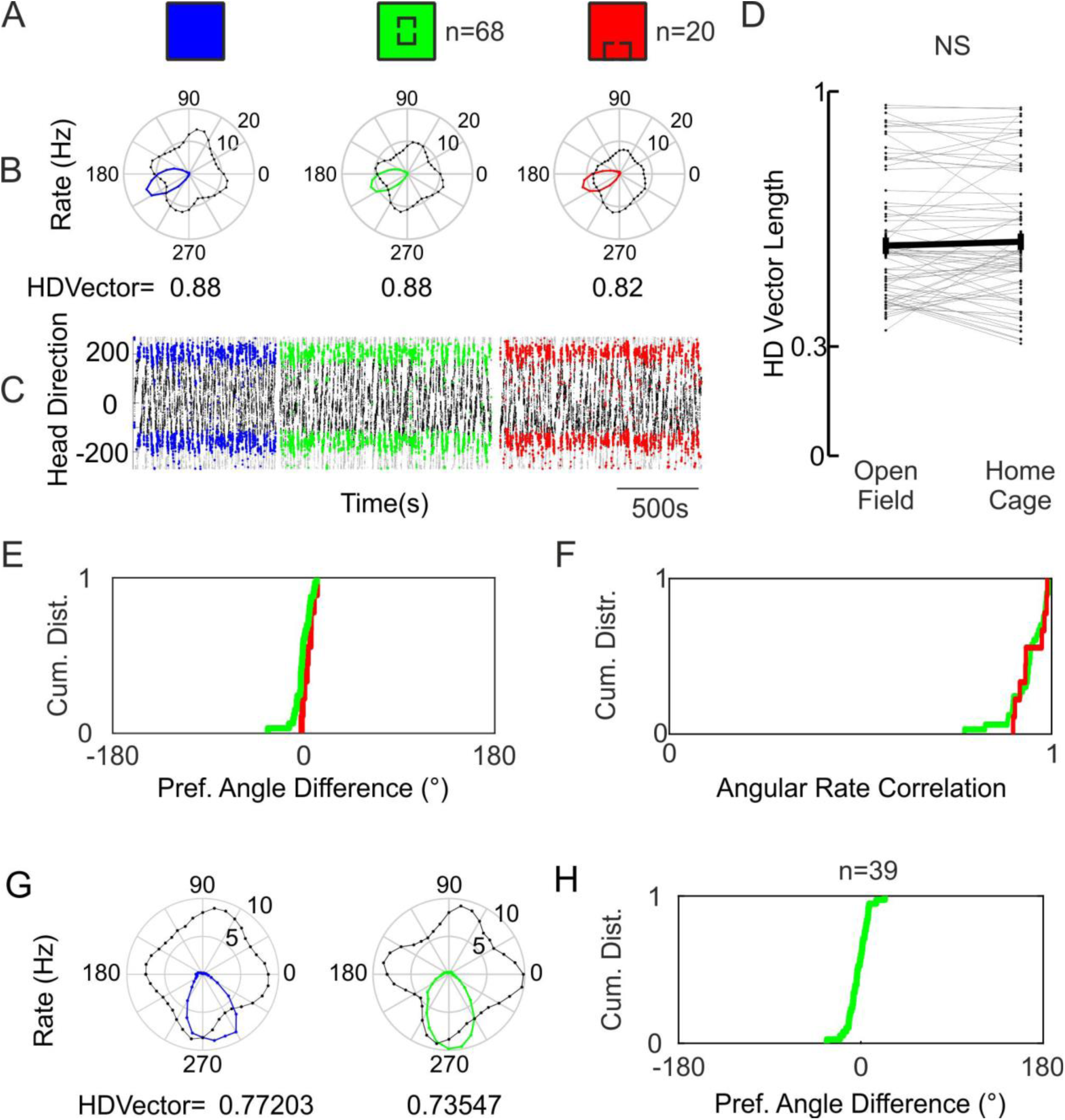
Head direction discharge is not affected by home cage location. A) Schematic of the conditions depicted for a HD cell, Open Field (Blue), Home Center (green), Home Moved (Red). B) Head direction rate polar plots (rate in corresponding color, angular occupancy in black). C) Spikes of the HD cell are clearly preferring a stable head direction. Black and grey, Head direction of the animal in time (duplicated for visualization). Spikes are plotted on top in the corresponding RGB color scheme. D) Head Direction Rayleigh vector lengths are not affected by the presence of the home cage. (KS Normality test, p=0.34, t-test, p=0.96, N=68) E) Cumulative frequency function of the distributions of differences between preferred angle in the Open Field and in each Home Condition. Note that the inflection point is at 0 and the very steep slope. (KS Normality Test for Angle Difference, p=0.6993) F) Correlation between the Angular Rates of both home conditions in relation to the open field. Cumulative frequency graph shows that correlations are distributed close to 1. Black lines represent individual cells. Colored lines represent the mean and SE for each condition. G) Head direction polar plots of a conjunctive grid cell show a consistent head directionality between sessions: Open Field (blue) and home center (green) conditions. H) Cumulative distribution of Preferred Angle differences in conjunctive grid cells.. (KS Normality Test, p=0.26, N=39)

### Absence of explicit home vector cells

As a second step, we assessed if the home cage could induce an explicit home vector representation, i.e. neural discharges tuned to the direction of the home cage (Figure 3). We therefore computed home direction as schematized in Figure 3A, where a home direction of 0 corresponds to when the animal is facing the home (Figure 3A bottom). We performed this computation fictively in the open field in absence of home cage (blue Figure 3B top, relative to where the center of the home cage would later appear) and relative to the real home cage (green Figure 3B top). Differences between the results of these two computations could be indicative of a home vector representation. In Figure 3C-D we show data from one of the non-grid and non-Head Direction cells with the strongest home direction tuning in terms of vector length. As shown in the polar rate plot and in the time resolved distribution of spikes in the home direction space (Figure 3B middle and bottom, respectively), even in this cell there is no strong home direction tuning (Vector Length=0.28). We found that in the general population of cells the distribution of Rayleigh vector for the resulting angular firing rates remains mostly below cut-off level used for similar variables like head directionality vector length (0.35 in our case; Figure 3E). We then investigated home direction tuning in both grid cell and head direction populations recorded in MEC and PaS to test for a representation of home direction in a conjunctive way. This analysis did not reveal any home direction tuning. Figure 3F-G shows an example corresponding to a head direction cell with a very strong head direction vector (Figure 3F) and a very weak home vector (Figure3G). We compared home direction vectors and Head Direction vectors of the HD cell population and found a large difference in their distribution strongly favoring Head Directionality (Figure 3H). Besides presenting very low Home Direction vectors, the presence of the home did not affect the encoding of home vectors in the HD cell population (Figure 3I). Similarly, grid cells did not represent home direction vectors. For the grid cell population we found very short home vector lengths and no effect of the introduction of the home in the environment (Figure 3K). These data indicate that there is no home vector tuning in the entorhinal and parasubicular cells.

**Figure 3.**
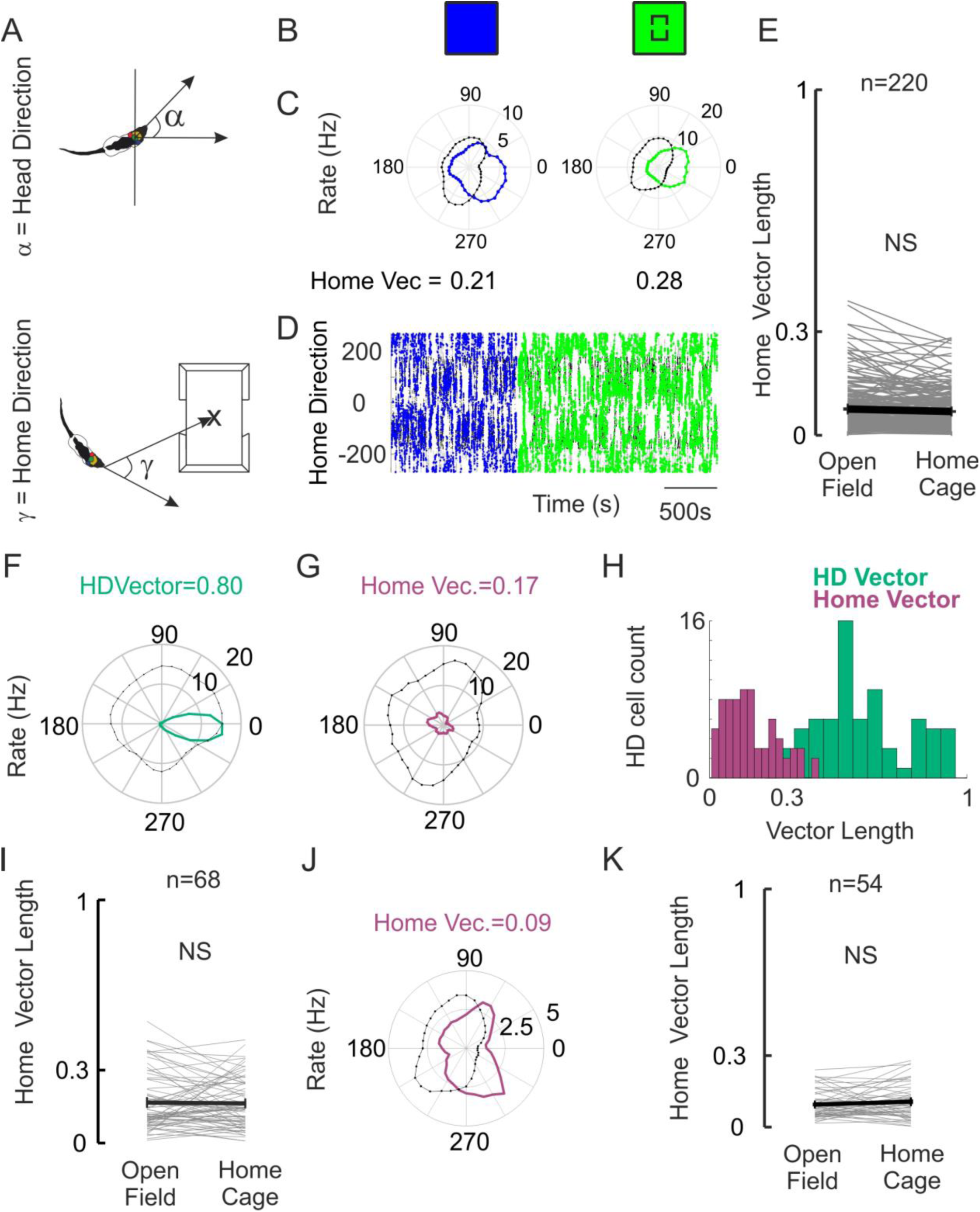
The home cage does not induce home vector discharges. A) Top: Representation of Head Direction Angle. Bottom: Representation of Home Direction angle. Note that in this case corresponds to the angle between the head direction vector and the vector pointing from the head of the animal to the center of the home(x). Hence, the animal facing the home retrieves a value of 0 and the animal running away from the home results in a value of 180 or −180 B) Schematic of the conditions depicted for a non-grid and non-HD cell, Open Field (Blue), Home Center (green) C) Polar Plots of Home Direction Rates (blue and green according to B, showing no clear Home Direction vectors). D) Plot of the Home Directionality of the spikes for this cell, shows no clear home direction preference. E) Distribution of fictive Home Direction Vector Lengths in the Open Field (calculated to the center of the arena) and the corresponding vector lengths once the home cage is placed at the center of the arena. Mean and SE are depicted in black showing no significant increase in vector length. (KS Normality, p=0.0037, Signed-Rank Test, p=0.21, N=220) F). The head direction vector of one Head Direction cell. G) Note the lack of clear home vector tuning curve for the same cell as in F. H) The distribution for all pure HD cells of Home Vector Lengths is much smaller than for head direction. I) Home vector lengths of pure HD cells did not change with the presence of the home. (KS Normality, p=0.367, t-test, p=0.560, N=68) J) Home Direction polar plot for pure Grid cell, showing lack of home vector. J K) Home Vector Lengths for all Grid cells are very low and did not change with the presence of the home. (KS Normality, p=0.780, t-test, p=0.250, N=54)

### Globally stable grid cells translocate single fields towards the home cage

Thirdly, we assessed how the home cage affects positional signals and in particular grid cell discharges (Figure 4). Already implied by the stability of the head direction system to home cage insertion, the animal must remain familiar with the global environment. In line with the lack of effect on head direction cells, grid cells did not globally remap because of home cage insertion. Global average firing rates were not altered by the presence of the home (KS Normality, p=0.384, ttest, p=0.541, N=51). We observed, however, that a proportion (60%) of grid cells recorded in both the parasubiculum and the medial MEC in the presence of the home cage did alter their discharge patterns. Figure 4A-B shows a grid cell recorded in the MEC for which the introduction of the home in the center of the arena retained the global representation but resulted in a local shift of a single firing field towards the home. It can be clearly observed how one central field in the green Home Cage in the center condition is shifted towards the position of the Home Cage. This is visible both at the level of spike positions (Figure 4B Top) where the green spikes are clearly shifted in position or in Figure 4B Bottom, where a composite normalized rate map using the RGB color scheme also clearly depicts the displacement of the central grid field in the green condition. Figure 4C presents two further examples of grid cells modifying their activity by translocating fields towards the location of the home-cage. We wondered if these shifts affect rates of grid cells inside the home cage. In order to be able to compare grid cells with distinct firing rates, and phases, we looked at rates of individual cells normalized to the average rate of the cell. If we compare normalized rates of the grid cells inside the home, with the normalized rates in the equivalent area of space during the open field session, we note a significant increase in these rates in the population (Figure 4D). This change is also evident in the change in the profile of normalized rates with the Euclidian distance to the center of the home (Figure 4E). Spatially averaging peak normalized rate maps of all grid cells in the open field and in the home center condition shows a local spatial increase in the firing rate, which can be further visualized by calculating the difference between the averages (Figure 4F-right). Similar results can be obtained by normalization of spatial maps to the mean (results not shown). However, not all grid cells increased their normalized rate inside the home, a smaller fraction did not up-modulate its firing rate (Figure 4G). We used the unity line as an *ad hoc* classification to disentangle possible differences due to the original configuration of the grid. Once we separated these two populations and performed the spatial averages, and euclidian average (Figure 4H) it became clear that positively modulating cells contribute to the overall effect. This result is of course tautological, but we observed in addition, that the average of these cells in the open field session had a low rate in the area where the home was to be placed originally. These observations indicate that introducing the home cage boosted firing in cells, via node translocation, that did initially had no grid firing node at the position where the Home Cage would be placed (Figure 4H Top). On the other hand, unchanged cells, tend to have an original field in the center of the arena, where the home will be placed (Figure 4H Bottom). Collectively, these observations show that the grid cells, which do not have a firing node in the home location, locally alter their firing patterns. Specifically it appears that a firing nodes close to the home cage get ‘sucked’ into home location.

**Figure 4.**
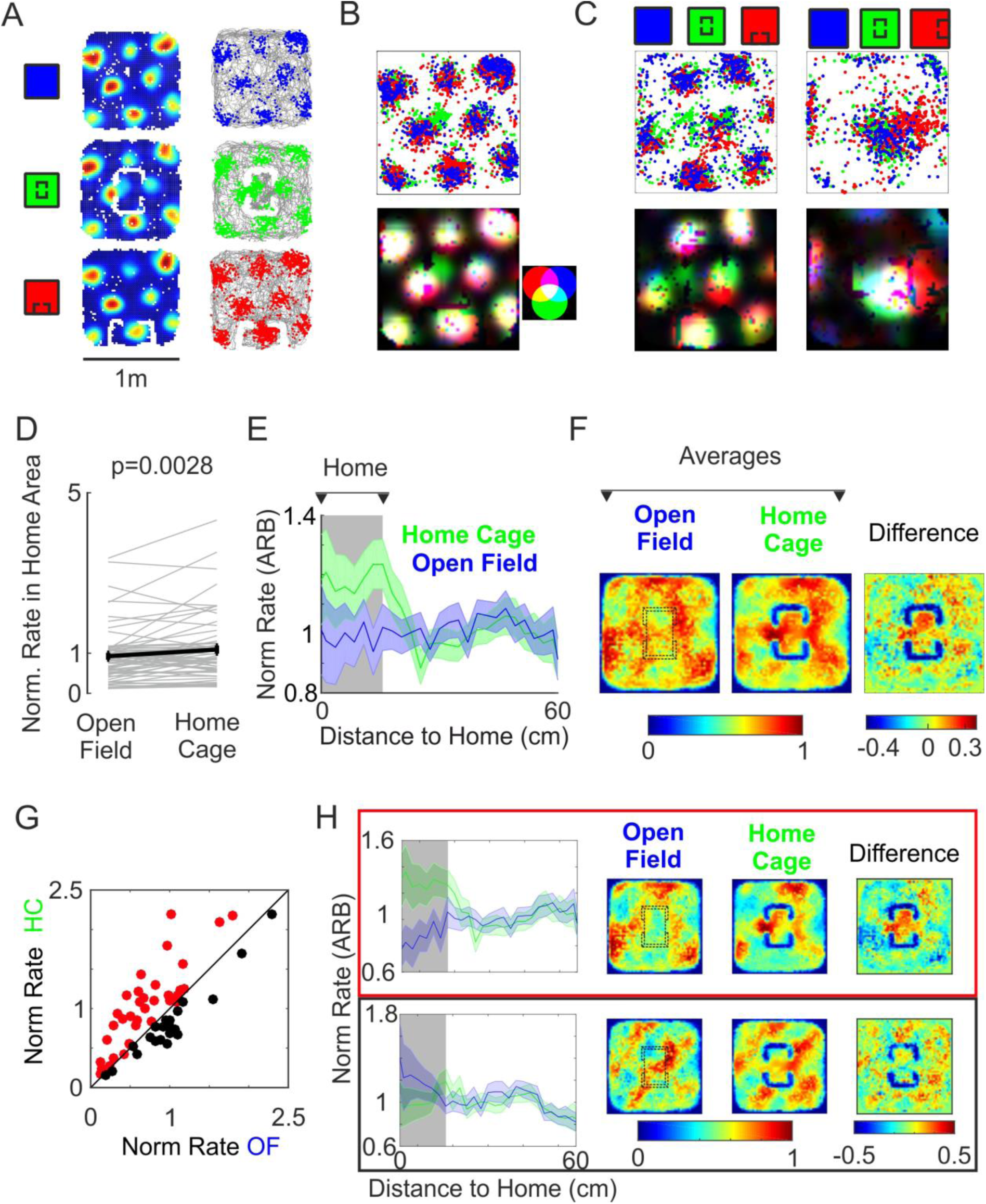
Single Firing Fields of Grid Cells shift towards Home Cage Location. A) Grid Cell under 3 conditions, Open Field, Home Center and Home Moved (Blue, Green, Red respectively). Left: Normalized Rate Maps. Right: Spikes (RGB) superimposed on the rat trajectory (grey). B) Composite plot of the Spike Positions for the three conditions of the cell in A. Note the change in positions for Home Center spikes. Bottom; Composite Rate Map, each rate map is normalized and assigned the corresponding channel in an RGB image. Side Panel demonstrates the color compositions for the different RGB mixtures. (Cell Recorded in dorsal medial MEC) C) Two parasubicular Grid Cells, under the same 3 conditions as in B. Note how single fields move towards the position of the home. D) Normalized firing rate increases (ARB: arbitrary units) in a region of the arena corresponding to the location of the home in comparison to the same area in the open field condition (KS Normality, p=0.280, t-test, p=0.0029, N=51). E) The increase in normalized rate is evident in the Euclidian profile to the center of the home. Solid line corresponds to mean normalized rate, and shaded area correspond to the SEM. Green: Home center. Blue: Open Field. F) The same effect is evident in the spatial averages of the peak normalized rates. Spatial Averages of peak normalized rate maps for all cells in the Open Field (Left) and Home Center (center). On the right the difference between these two average maps is shown G) Brute force split between cells up-modulating their normalized firing rate in the home area (Red, n=32) and not-up-modulating (Black. n=19) H) Top: Same scheme as F) for up-modulating cells from (G), with the inclusion of the Euclidian profile to the left. Bottom: Same scheme for down-modulating cells from (G), with the inclusion of the Euclidian profile to the left.

To quantify the local grid cell changes we performed a sliding window correlation analysis (Wernle et al., 2018) between the peak normalized spatial rate maps in the open field and home presence conditions. For any given pixel in the arena we selected a surrounding squared region (of dimension similar to the cells average grid spacing). This region matches spatially for both conditions (Figure 5A, boxes in white dotted lines). For each region the cell’s firing rates were correlated between Open Field and Home conditions. The result is a heat-map of local correlation values between the different conditions. In this example, the presence of the home reduced locally the correlation of the grid (Figure 5A, right). We averaged these correlation maps for all grid cells and found that these show a mean local decrease in correlation corresponding to the location of the home in the center (Figure 5B left). Notably, the decrease in mean correlation follows the home, when we moved it to the side of the arena (Figure 5B, right). The decorrelation of grid cell activity appears to be local and related to the position of the home in the environment. This analysis reinforces the idea that presence of the home resulted in a change in the local activity of the grid while not affecting the global encoding of space. We were curious whether this effect might be driven or intensified by the positive valence of the home or if grid cells are just encoding for the changes in local geometry introduced with the home. To disentangle this we performed additional controls with a cardboard box with equal dimensions to the home, yet without both the familiarity and the social valence of the home. We found, however that the correlated activity of grid cells in the presence of the box drove a similar local change with respect to the open field as had the home (Figure 5C Left). At the same time activity in the presence of the box correlated strongly with the activity in the presence of the home (Figure 5C Right). Hence, we believe that the effect seen of grid cells is indeed related to the change in the internal structure of the environment. This could be related to, just the presence of an “object” in the environment. To dissect this further we tested whether tall objects, which do not alter the space substantially, have a similar local effect on grid cells. We found that the simple presence of an object does not decrease the correlation of grids around it (Figure 5D). These results point towards the requirement of a change in the internal structure of the environment which would considerably affect the trajectories available to the rat. We wondered if single tall objects do not produce the same effect, perhaps because they minimally affect the availability of paths. Pursuing this idea further we presented the animals with new internal structures inside the environment. We used an open corridor of 70 × 10 cm, which we could place in the arena in different orientations. The presence of a corridor inside an open field grossly changes the availability of paths in the same familiar environment. Our prediction would be that grid cells would care about the position of this corridor and modify their firing fields to better represent this salient path. Introducing such corridor revealed big shifts in grid fields towards the corridor in some, but not all conditions (Figure 5E, moving field highlighted in pink). This example further demonstrates the effect of internal geometry in purely local grid cell activity.

**Figure 5.**
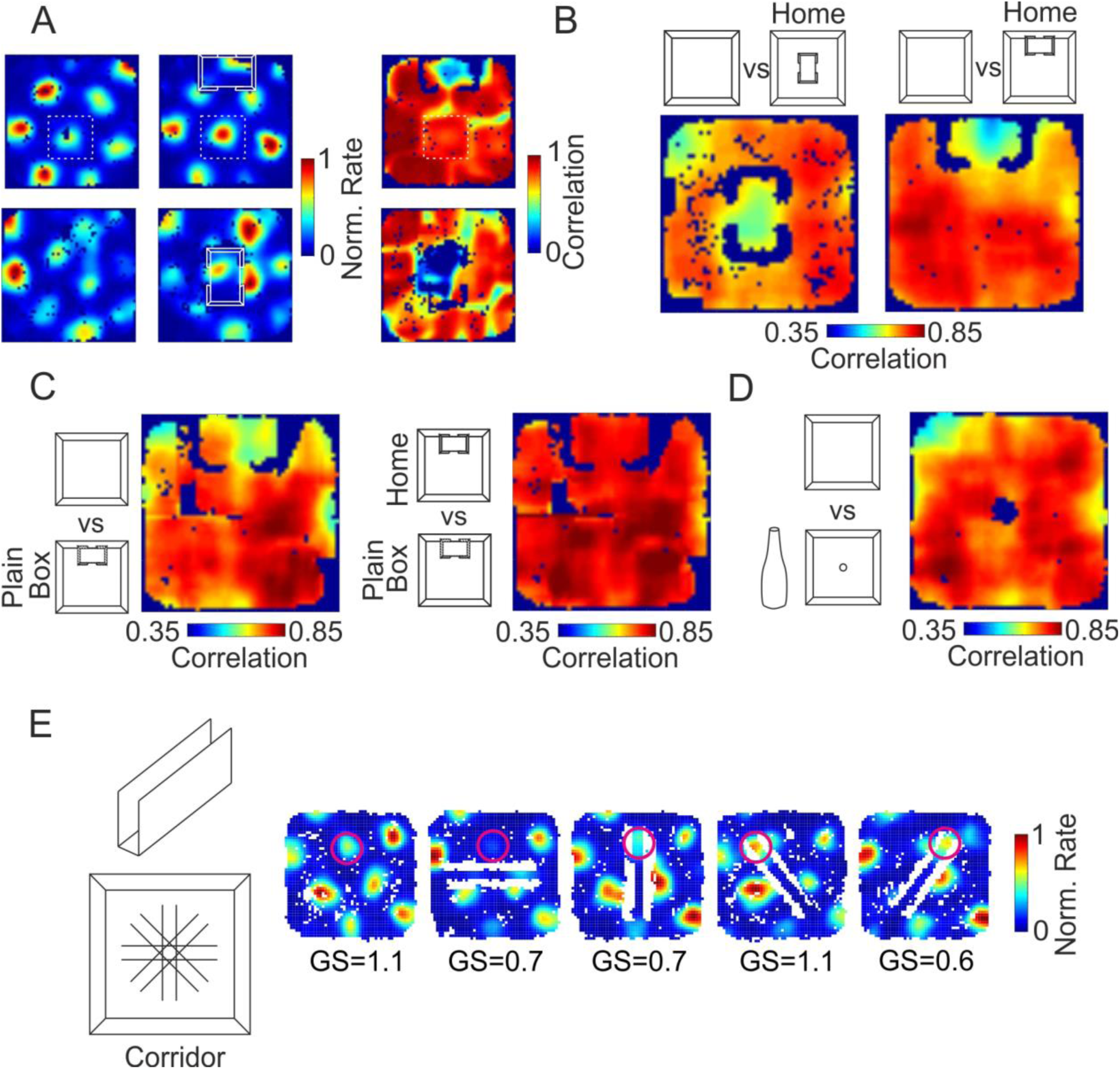
Effect of Home on grid cell population is driven by geometry. A) Sliding window correlation analysis between conditions. Top: Example of peak normalized rate maps of a Grid Cell in the Open Field (left) and Home North (center) condition, and the corresponding sliding window correlation between these two (right). Bottom: Same as top but for another cell compared to the Home Center condition. B) Sliding window correlation analysis for all Grid cells in both conditions. Note the lower average local correlation in the position of the home for both cases. C) Sliding window correlation analysis comparing the Home vs a cardboard box with similar dimensions, showing that the effect is related to geometry not valence of the home. Left: the comparison between the Open Field and the Cardboard box is shown. Right: High correlation comparison between the Home and the cardboard box. D) Sliding window correlation analysis between the Open Field and a Tall Object in the center of the arena did not produce low correlations near the object. E) The presence of a linear corridor in the arena in different angles produced shifts of grid fields in some of the conditions further demonstrating the strong effect of internal geometry. Moving field highlighted in pink.

## Discussion

The home-cage is a highly relevant location for the rat. Consistent with this idea the home cage was capable of inducing characteristics of natural homing behavior. Even though, the parasubiculum and the MEC are causally linked to the expression of spatial navigation, and contain a panoply of cells encoding variables linked to space and orientation, we did not find an explicit firing rate representation of a home vector. Specifically, in the same quality of needle like encoding as the goal direction cells found by Sarel et al in the hippocampus of freely flying bats with respect to a home-like platform (Sarel et al., 2017).

We found that both head direction cells and the head directional component of conjunctive grid cells are not affected by the presence of the home. The preferred head direction is retained, even if the home is translated to one of the sides of the arena breaking symmetry even further. This points to head direction cells and grid cells maintaining their encoding of global familiar environment, i.e., there is no remapping because of the appearance of the home. The fact that the rat was never removed from the arena or disoriented between sessions probably explains the consistency of the encoding for the global environment. Many of the traditional cue card rotation experiments, where head direction realignment was found were performed in cue-deprived conditions and with a step of disorientation of the animal between sessions.

Even though earlier work had shown that grid cell macro crystalline hexagonal patterns break in complex shaped environments like the hairpin maze, and more recent studies have shown global distortions are possible when exposing the animal to irregular shaped environments or changing external boundaries (Krupic et al., 2015, 2018), we are just beginning to understand how the structure of the environment affects single firing fields and whether these distortions might contribute encoding more intricate environments.

In similar manner a study by Wernle and collaborators shows a much faster change in the grid pattern of cells encoding two environments separated by a wall, before and after said wall is removed (Wernle et al., 2018). Grid cells showed a tendency towards a more global representation by stitching up the previous two global grid patterns, into a meta-global grid pattern. This intriguing study points to a strong role of borders in the establishment of the grid pattern. However, in this case it is hard to disentangle what might be the single contribution of local structures, since the overall global environment is changed in shape and size.

Part of our contribution lies in testing what are the micro level effects of internal structures of the environment (in our case the rat’s home) on grid fields of well learnt, familiar environments without disturbing the global environment itself. In line with the head direction, grid cells did not globally remap with the presence of the home. In all cases they retained their previous phase, orientation and spacing. However, we did find that grid cells are far from being a perfect crystal projected on to the arena. We observed shifts of local firing fields towards the position of the home. This phenomenon manifested itself by a local decrease in correlation of the rate maps of the grid cells in the vicinity of the home, while global correlation of the grids is preserved. This result falls in line with recent work showing that single grid fields moved to reward locations (Boccara et al., 2019). However, we did not see any global remapping (Butler et al., 2019) of head direction cells or grid cells due to the simple presence of the home, even though we do see an increase in the home location representation because of field convergence.

The relocation of grid fields due to changes in the internal structure of the environment may have a critical role in vector based navigation and route planning. The most recent interpretation of the role of grid cells in spatial navigation relates to the possibility of using grid populations to directly calculate the shortest path between two points (Kubie and Fenton, 2009; Banino et al., 2018). Changes in the internal structure of the environment restructure the availability of paths in the environment. Some paths need to be better represented, while some others are now impossible. The hexagonal grid cell in the open field, might be the best way of encoding all possible segments, which for example includes all possible directions. As has been described, when the same cells are recorded in a hairpin maze, grid cells become linearized inside each transect and repeating for different pins of the maze and at the same time becomes dependent on the direction of travel. This restructuring of the grids is in line with the restructuring of possible paths in the arena, the only vectors that can be calculated are ones going back and forth through each hairpin. When rats were trained to perform a zig-zag in an open field environment, behaviorally mimicking the hairpin like behavior, the gridness of the cells were once again present, given that all the possible paths were again available (Derdikman et al., 2009). A similar reasoning can be applied to recent results showing grid cells preferentially encoding memorized goal locations (Boccara et al., 2019; Butler et al., 2019). A goal location changes the affordance of space and the type of paths in space that are to be encoded.

The shifts found for grid cells together with the up-modulation of firing rates in the home point towards an increased overlap in the grid population firing in the position of the home. This could allow for a more resolved encoding of that area of the environment. Our results are in-line with recent work showing the adaptability of grid cells to changes to their global environment (Krupic et al., 2018; Wernle et al., 2018). In addition, we have shown that grid cells also flexibly encode local internal changes in the environment, pointing towards their role in encoding more naturalistic environments and suggesting a clear hypothesis towards their role in allowing vectorial navigation in complex environments.

While our study confirms on a behavioral level that the home cage is a unique location of rats, we have not been able to decipher a neural signature of what makes home a special place. We wonder, if such a ‘neural home signature’ exists in the cortico-hippocampal system or if subcortical circuits provide the animal with this information.

## Materials and methods

All experimental procedures were performed according to the German guidelines on animal welfare under the supervision of local ethics committees.

### Subjects

We obtained data from 6 male Long Evans rats (~300g) using chronic extracellular recordings.

### Tetrode recordings

Tetrode recordings from parasubiculum and MEC were performed, as recently described (Tang et al., 2016).

Tetrodes were turned from 12.5 μm diameter nichrome wire (California Fine Wire Company) and gold plated to 250-300 kΩ impedance. In order to identify tetrodes in the complex anatomy of the Parasubiculum and MEC, tetrodes we stained with fluorescent tracers DiI and DiD (ThermoFisher, Scientific) before implantation.

Rats were chronically implanted with 32 channel Harlan-8 Drives (Neuralynx) with independently movable tetrodes.

Spiking activity and local field potential were recorded at 32 kHz (Neuralynx; Digital Lynx) or at 32 kHz using the wire free RatLogger-32 from Deuteron Technologies Ltd. All recordings were done freely-moving during behavioral tasks. The animal’s location and head direction was automatically tracked at 25 Hz by video tracking with a colored camera using Red-Blue head-mounted LEDs or Red-Blue colored plastic targets. After recordings, the animals were transcardially perfused. Spikes were detected and clustered using Kilosort (Pachitariu et al., 2016) and manually curated based on PCA overlap and refractory period violations using the visualization toolbox Phy.

### Behavioral Procedures

After surgery animals were adapted for two weeks to a modified home-cage with 2 side doors. Concurrently, the animals were put under food restriction up to achieving 80% of their adlibitum feeding body weight, and adapted to foraging small chocolate treats while familiarized with a cue rich 1×1m arena under well-lit conditions.

Once the tetrode reached the parasubiculum and MEC (based on the presence of strong theta), we recorded from 2 to 8 sessions per day of 12-18 minute each, while the animal explored the same familiar arena without removing or disorienting the rat between sessions. In between each session we introduced, or displaced the home-cage of the animal or additional control objects (Bottle, Plain Box, and Corridor).

### Hoarding

In a subset of sessions (n=5 rats) we performed hoarding behavioral tests. For these we positioned the home-cage in the center of the arena, and instead of randomly dispersing chocolate treats, we dispersed standard food pellets outside the rats home-cage. Food deprived rats, retrieved these pellets and horded them inside the home cage without any specific training. Rat’s hoarded up to 80 pellets in 20 minutes.

### Hoarding Task vs No Task

To dissociate the possible effect of the home location with the effect of the behavioral task, neural recordings where perform comparing ‘No Task’ behavior. That is to say, that both in absence (open field) or presence of the home, rats were simply randomly foraging for minimal sugary treats. This allowed for a fair behavioral comparison and the necessary occupancy for grid cell analysis.

### Histology

After perfusion, the brain was post-fixed in PFA 4% for 12-18hrs. The brain was then sectioned tangentially using the methods described in (Lauer et al., 2018) and recording sites assigned by histology using Inmunohistochemistry of Calbindin to correctly assign the parasubiculum and MEC recordings.

We did not see significant differences in the populations and pooled cells from parasubiculum, and MEC.

### Analysis of Spatial Modulation

The position of the rat was defined as the midpoint between two head-mounted LEDs or colored targets. A running speed threshold (of 5cm/s) was applied for isolating periods of rest from active movement. Color-coded firing maps were plotted. For these, space was discretized into pixels of 2 cm × 2 cm, for which the occupancy z of a given pixel × was calculated as

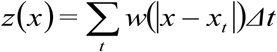

where *x*_*t*_ is the position of the rat at time *t, Δt* the inter-frame interval, and w a Gaussian smoothing kernel with σ = 5cm.

Then, the firing rate r was calculated as

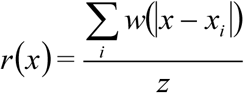

where *x*_*i*_ is the position of the rat when spike *i* was fired. The firing rate of pixels, whose occupancy *z* was less than 20 ms, was considered unreliable and not shown.

For spatial and head-directional analysis, both a spatial (> 50% spatial coverage) and a firing rate inclusion criterion (> 0.5 Hz) were applied. Spatial coverage was defined as the fraction of visited pixels (bins) in the arena to the total pixels.

### Analysis of Spatial Information

For all neurons, we calculated the spatial information rate, *I*, from the spike train and rat trajectory:

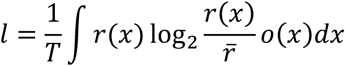

where *r(x)* and *o(x)* are the firing rate and occupancy as a function of a given pixel × in the rate map. 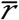 is the overall mean firing rate of the cell and *T* is the total duration of a recording session (Skaggs et al., 1993). A cell was determined to have a significant amount of spatial information, if the observed spatial information rate exceeded the 95th percentile of a distribution of values of I obtained by circular shuffling. Shuffling was performed by a circular time-shift of the recorded spike train relative to the rat trajectory by a random time for 1000 permutations.

### Analysis of Grid Cells

Analysis of Gridness. Grid scores were calculated, using publically available codes from the Derdikman Lab’s recent publication (Ismakov et al., 2017). By taking the autocorrelogram, centered on, but excluding the central peak. The Pearson correlation of the autocorrelogram with its rotation for 60 degrees and 120 degrees was obtained (on peak rotations) and also for rotations of 30 degrees, 90 degrees and 150 degrees (off peak rotations). Gridness was defined as the minimum difference between the on-peak rotations and off-peak rotations.

### Analysis of head directionality

Head-direction tuning was measured as the eccentricity of the circular distribution of firing rates. For this, firing rate was binned as a function of head-direction (n = 36 bins). A cell was said to have a significant head-direction tuning, if the length of the average vector exceeded the 95th percentile of a distribution of average vector lengths calculated from shuffled data and had a Rayleigh vector length > 0.35. Data was shuffled by applying a random circular time-shift to the recorded spike train for 1000 permutations.

We studied the head directional properties across subsequent conditions with the presence of the home by analyzing changes in both the angle of the Rayleigh vector or the modulus of the vector.

### Home Direction Analysis

Home Direction is calculated as in Figure 3A, as the angle the rat would have to turn its head to face in the direction of the home. In other words the angle between 2 vectors defined by the head direction of the animal and a vector between the position of the animal and the position of the home. We applied the same Rayleigh vector analysis to cells in the Home Direction space, in this case finding no significant cells.

### Classification of cells into functional categories

Cells were classified as head-direction cells, pure grid cells, conjunctive grid cells and rest cells, based on their grid score, spatial in-formation and significance of head directionality according to the following criteria:

- Head direction cells: Rayleigh Head Direction vector length > 0.3 & significant head-direction tuning
- Pure Grid cells: Grid score > 0.42 & significant spatial information.
- Conjunctive Grid Cells: pass criteria for both Grid Cells and Head Direction cells.
- All other cells are consider in the Rest category.

Besides the cell classification, for comparison of different sessions for the same cells we used a stability criterion for cell removal if the Z-score of the firing rate for a session goes below

- 1.5 of the previous sessions.

### Grid cell sliding window correlation analysis

Given the overall global stability of grid cells, in order to compare the population of grid cells in the presence or absence of the home we used a sliding window correlation method as described in Wernle et al (Wernle et al., 2018).

For each cell we start with two normalized rate maps of the cell, one for each condition being compared. We choose a fixed window of size corresponding to the spacing of the grid cell. Starting in one corner of the rate maps, we correlate between both maps the rate of active bins covered by the same window. We assign this local correlation value to the position of the center bin of such window. By sliding the window along all the bins of the rate maps we end up with a local correlation heat map that allows us to dissect local changes in each grid cell resulting from the introduction of the home. In all cases, non-existent values were removed from the correlations.

To compare the effect of the home in the population we averaged the local correlation heat maps.

## Acknowledgements

This study was funded by the Einsteinstiftung (ESB project ‘Dynamics of electronically coupled neuronal networks” A-2016-350). We acknowledge the outstanding technical assistance of Undine Schneeweiss and Juliane Steger. We thank lab members S.R., E.C., J.S., for discussions regarding the project and proofreading of the manuscript. We also extend thanks to members and alumni of TENSS.

## Competing Interests

We declare no competing interests.

## References

Alleva E, Baldaccini NE, Foà A, Visalberghi E (1975) Homing Behaviour of the Rock Pigeon. Monit Zool Ital - Ital J Zool 9:213–224.

Banino A et al. (2018) Vector-based navigation using grid-like representations in artificial agents. Nature:1.

Boccara CN, Nardin M, Stella F, O’Neill J, Csicsvari J (2019) The entorhinal cognitive map is attracted to goals. Science 363:1443–1447.

Boccara CN, Sargolini F, Thoresen VH, Solstad T, Witter MP, Moser EI, Moser M-B (2010) Grid cells in pre- and parasubiculum. Nat Neurosci 13:987–994.

Butler WN, Hardcastle K, Giocomo LM (2019) Remembered reward locations restructure entorhinal spatial maps. Science 363:1447–1452.

Derdikman D, Whitlock JR, Tsao A, Fyhn M, Hafting T, Moser M-B, Moser EI (2009) Fragmentation of grid cell maps in a multicompartment environment. Nat Neurosci 12:1325–1332.

Diehl GW, Hon OJ, Leutgeb S, Leutgeb JK (2017) Grid and Nongrid Cells in Medial Entorhinal Cortex Represent Spatial Location and Environmental Features with Complementary Coding Schemes. Neuron 94:83–92.e6.

Fyhn M (2004) Spatial Representation in the Entorhinal Cortex. Science 305:1258–1264.

Gill RE, Piersma T, Hufford G, Servranckx R, Riegen A (2005) Crossing the ultimate ecological barrier: evidence for an 11 000-km-long nonstop flight from alaska to new zealand and eastern australia by bar-tailed godwits. The Condor 107:1–20.

Hafting T, Fyhn M, Molden S, Moser M-B, Moser EI (2005) Microstructure of a spatial map in the entorhinal cortex. Nature 436:801–806.

Ismakov R, Barak O, Jeffery K, Derdikman D (2017) Grid Cells Encode Local Positional Information. Curr Biol 0 Available at: http://www.cell.com/current-biology/abstract/S0960-9822(17)30771-6 [Accessed July 28, 2017].

Krupic J, Bauza M, Burton S, Barry C, O’Keefe J (2015) Grid cell symmetry is shaped by environmental geometry. Nature 518:232–235.

Krupic J, Bauza M, Burton S, O’Keefe J (2018) Local transformations of the hippocampal cognitive map. Science 359:1143–1146.

Kubie JL, Fenton AA (2009) Heading-vector navigation based on head-direction cells and path integration. Hippocampus 19:456–479.

Lauer SM, Schneeweiß U, Brecht M, Ray S (2018) Visualization of Cortical Modules in Flattened Mammalian Cortices. JoVE J Vis Exp:e56992–e56992.

Maaswinkel H, Whishaw IQ (1999) Homing with locale, taxon, and dead reckoning strategies by foraging rats: sensory hierarchy in spatial navigation. Behav Brain Res 99:143–152.

Menzel R, Greggers U, Smith A, Berger S, Brandt R, Brunke S, Bundrock G, Hülse S, Plümpe T, Schaupp F, Schüttler E, Stach S, Stindt J, Stollhoff N, Watzl S (2005) Honey bees navigate according to a map-like spatial memory. Proc Natl Acad Sci 102:3040–3045.

Mittelstaedt ML, Mittelstaedt H (1980) Homing by path integration in a mammal. Naturwissenschaften 67:566–567.

Neave F (1964) Ocean Migrations of Pacific Salmon. J Fish Res Board Can 21:1227–1244.

Pachitariu M, Steinmetz NA, Kadir SN, Carandini M, Harris KD (2016) Fast and accurate spike sorting of high-channel count probes with KiloSort. In: Advances in Neural Information Processing Systems 29 (Lee DD, Sugiyama M, Luxburg UV, Guyon I, Garnett R, eds), pp 4448–4456. Curran Associates, Inc. Available at: http://papers.nips.cc/paper/6326-fast-and-accurate-spike-sorting-of-high-channel-count-probes-with-kilosort.pdf.

Ray S, Burgalossi A, Brecht M, Naumann RK (2017) Complementary Modular Microcircuits of the Rat Medial Entorhinal Cortex. Front Syst Neurosci 11 Available at: https://www.ncbi.nlm.nih.gov/pmc/articles/PMC5385340/.

Sarel A, Finkelstein A, Las L, Ulanovsky N (2017) Vectorial representation of spatial goals in the hippocampus of bats. Science 355:176–180.

Skaggs WE, McNaughton BL, Gothard KM (1993) An Information-Theoretic Approach to Deciphering the Hippocampal Code. In: Advances in Neural Information Processing Systems 5 (Hanson SJ, Cowan JD, Giles CL, eds), pp 1030–1037. Morgan-Kaufmann. Available at: http://papers.nips.cc/paper/671-an-information-theoretic-approach-to-deciphering-the-hippocampal-code.pdf.

Stensola T, Stensola H, Moser M-B, Moser EI (2015) Shearing-induced asymmetry in entorhinal grid cells. Nature 518:207–212.

Tang Q, Burgalossi A, Ebbesen CL, Sanguinetti-Scheck JI, Schmidt H, Tukker JJ, Naumann R, Ray S, Preston-Ferrer P, Schmitz D, Brecht M (2016) Functional Architecture of the Rat Parasubiculum. J Neurosci 36:2289–2301.

Taube JS (1995) Head direction cells recorded in the anterior thalamic nuclei of freely moving rats. J Neurosci 15:70–86.

Taube JS, Muller RU, Ranck JB (1990) Head-direction cells recorded from the postsubiculum in freely moving rats. I. Description and quantitative analysis. J Neurosci 10:420–435.

Tchernichovski O, Benjamini Y, Golani I (1998) The dynamics of long-term exploration in the rat. Biol Cybern 78:423–432.

Tsoar A, Nathan R, Bartan Y, Vyssotski A, Dell’Omo G, Ulanovsky N (2011) Large-scale navigational map in a mammal. Proc Natl Acad Sci 108:E718–E724.

Valerio S, Taube JS (2012) Path integration: how the head direction signal maintains and corrects spatial orientation. Nat Neurosci 15:1445–1453.

Wallace DG, Martin MM, Winter SS (2008) Fractionating dead reckoning: role of the compass, odometer, logbook, and home base establishment in spatial orientation. Naturwissenschaften 95:1011–1026.

Wernle T, Waaga T, Mørreaunet M, Treves A, Moser M-B, Moser EI (2018) Integration of grid maps in merged environments. Nat Neurosci 21:92.

Whishaw IQ, Gharbawie OA, Clark BJ, Lehmann H (2006) The exploratory behavior of rats in an open environment optimizes security. Behav Brain Res 171:230–239.

Winter SS, Blankenship PA, Mehlman ML (2018) Homeward bound: The capacity of the food hoarding task to assess complex cognitive processes. Learn Motiv 61:16–31.

